# Untapped microbial composition along a horizontal oxygen gradient in a Costa Rican volcanic influenced acid rock drainage system

**DOI:** 10.1101/663633

**Authors:** Alejandro Arce-Rodríguez, Fernando Puente-Sánchez, Roberto Avendaño, Eduardo Libby, Raúl Mora-Amador, Keilor Rojas-Jimenez, Dietmar H. Pieper, Max Chavarría

**Author notes:** Department of Molecular Bacteriology, Helmholtz Centre for Infection Research, 38124, Braunschweig, Germany (Alejandro Arce-Rodríguez). **Correspondence to:** Max Chavarría, Escuela de Química & Centro de Investigaciones en Productos Naturales (CIPRONA), Universidad de Costa Rica, Sede Central, San Pedro de Montes de Oca, San José, 11501-2060, Costa Rica.

## Abstract

Based on the analysis of 16S rRNA gene metabarcoding, here we report the shift in the microbial community structure along a horizontal oxygen gradient (0.40-6.06 mg L^−1^) in a volcanic influenced acid rock drainage (VARD) environment, known as San Cayetano (Cartago, Costa Rica; pH =2.94-3.06, sulfate ~0.87-1.19 g L^−1^, iron ~35-61 mg L^−1^). This VARD is dominated by microorganisms involved in the geochemical cycling of iron, sulfur and nitrogen; however, the identity of the species changes with the oxygen gradient along the river course. The anoxic spring of San Cayetano is dominated by a putative anaerobic sulfate-reducing Deltaproteobacterium as well as sulfur-oxidizing bacteria (such as *Acidithiobacillus* or *Sulfobacillus*), which favor the process of dissolution of sulfide minerals and oxidation of H2S. In oxic conditions, aerobic iron-oxidizers (*Leptospirillum, Acidithrix, Ferritrophicum, Ferrovum*) and heterotrophic bacteria (Burkholderiaceae Betaproteobacterium, *Trichococcus, Acidocella*) were identified among others. Thermoplasmatales archaea closely related to environmental phylotypes found in other ARD/AMD niches were also found throughout the entire ecosystem. This work describes the changes in bacterial diversity, and possible metabolic activities occurring along a horizontal oxygen gradient in a volcanic influenced acid rock drainage system.

## Introduction

Water bodies with elevated metal content and very high acidity are found throughout the world as the result of volcanic activity and/or biotic/abiotic processes as in acid rock drainage (ARD) sites. In ARD environments, water emerges from underground having a low pH value and remarkable amounts of chemical species such as iron, sulfate and other metals (Baker and Banfield 2003; Johnson *et al*. 2009; Sánchez-Andrea *et al*. 2011; Arce-Rodríguez *et al*. 2019). Most of the studied ARD ecosystems in the world have been caused by human activity, especially mining processes. The anthropogenic influence modifies the geology, hydrology and biology of the ARD, making difficult to pinpoint the inherent chemical and natural microbiological conditions of these sites. It is therefore of great interest to study ARD ecosystems with no history of anthropogenic influence (*i.e*. no mining). Some works on the chemistry and microbiology of natural ARD systems include the Pastoruri Glacier area in Huascarán National Park (Perú) (González-Toril *et al*. 2015), the Antarctic landmass (Dold *et al*. 2013), the Río Sucio (Braulio Carrillo National Park, Costa Rica) (Arce-Rodríguez *et al*. 2017) and the Río Tinto, located at the Iberian Pyrite Belt (IPB) of Spain (González-Toril *et al*. 2003; López-Archilla *et al*. 2004; García-Moyano *et al*. 2012; Sánchez-Andrea *et al*. 2012). In the latter case, although the river and its source are located on a site that has been mined for thousands of years, it is believed by some authors that the conditions of extreme acidity and heavy metal pollution along the river are due to natural processes that were operative before mining in the region started five thousand years ago (Fernández-Remolar, Rodriguez and Gomez 2003; Amils, Fernández-Remolar and the IPBSL Team 2014). Due to its conditions, it is a good consensus that Rio Tinto should be considered a study model of ARD environments. On the other hand, the extreme acidity environments produced by discharge of the products of volcanic or hydrothermal activity (*e.g*. H_2_S oxidation or SO_2_ reaction with water) into water bodies like lakes and rivers, share most of their natural composition with that of ARD sites and are of interest because they can be unequivocally assigned to natural causes (Urbieta *et al*. 2015, Arce-Rodríguez *et al*. 2017). Thus, low pH values due to sulfuric acid and high metal content, especially iron, are a common factor to both ARD and volcanic influenced water bodies.

From a microbiological point of view, the common factor in natural ARD and volcanic environments is the presence of iron- and sulfur-oxidizing bacteria. For example, in the Pastoruri Glacier area the presence of *Acidithiobacillus*, a sulfur- and iron-oxidizing acidophilic bacterium, was detected (González-Toril *et al*. 2015). Dold *et al*. (2013) reported the formation of an ARD system in the Antarctic landmass due to the activity of psychrophilic acid mine drainage microorganisms found in cold climates, specifically *Acidithiobacillus ferrivorans* and *Thiobacillus plumbophilus*. In Costa Rica, a country with an important volcanic influence (Castellón *et al*. 2013) and tropical climatic conditions, Arce-Rodríguez *et al*. (2017) reported that Río Sucio (a pristine ARD), is dominated by *Gallionella* spp. and other chemolithoautotrophic iron- and sulfur-oxidizing bacteria. Studies in Rio Tinto revealed that eighty percent of the water column’s bacterial species corresponds to only three bacterial genera: *Leptospirillum, Acidithiobacillus* and *Acidiphilium*, all involved in iron cycling (González-Toril *et al*. 2003, 2010). Minor levels of other iron-oxidizing bacteria such as *Ferrimicrobium, Acidimicrobium* and *Ferroplasma* have also been detected in Rio Tinto.

In this work, we investigated the microbiological characteristics of the source and early course of an ARD riverine environment that has a contribution of sulfuric acid by volcanic activity as well as associated chemical species product of rock dissolution processes. This volcanic influenced acid rock drainage (VARD), known as San Cayetano Creek is located in a private farm northeast of Irazu volcano (Cartago, Costa Rica) (Fig. 1). Due to its location within a private property and difficult access this site has seen little human influence. This acid stream has a length of about 3 km from its source to its confluence with the previously described Rio Sucio (Arce-Rodríguez *et al*. 2017) but is only a minor tributary. Our results indicate that chemistry of San Cayetano is both the result of volcanic input and biological activity of iron- and sulfur-oxidizing bacteria distributed along a well-established oxygen gradient.

**Fig. 1.**
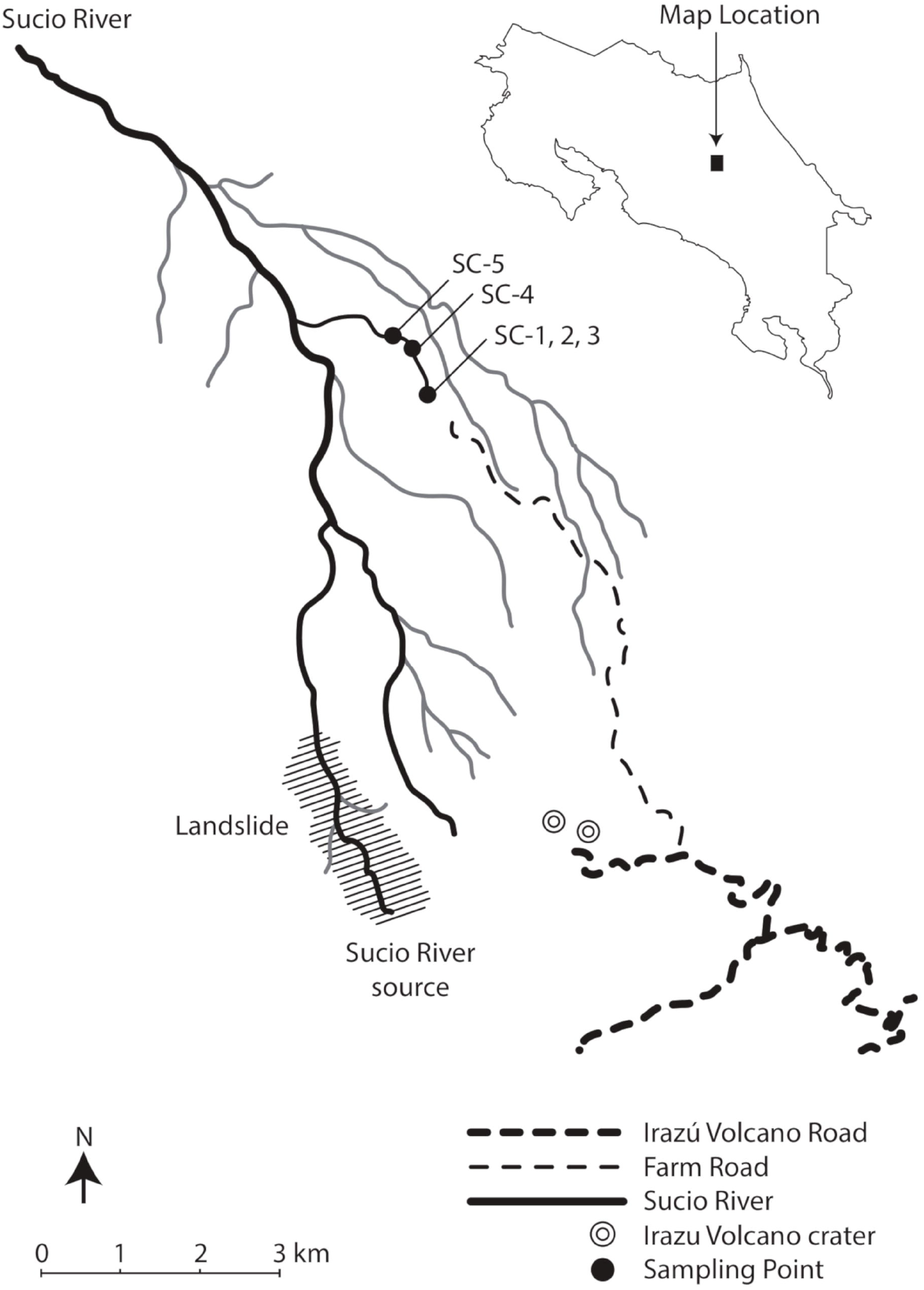
San Cayetano stream in Central Mountain Range, Costa Rica. The San Cayetano stream is located in the Central Valley, next to the Irazú volcano. This stream has a length of about 3 km to its mouth in the Rio Sucio.

## Materials and Methods

### Sampling and field measurements

The San Cayetano acid rock drainage system is located in a private farm in Cartago, Costa Rica. All the needed permits for sampling water and sediments were requested directly to the owners of the property. In August 2016, samples of water (three samples of 1 Liter each) were collected at different points along this stream (see Fig. 2): SC-1 (sampling point located at the origin, 10.032308 N 83.864449 W), SC-2 (10 meters from origin), SC-3 (30 meters from origin), SC-4 (600 meters from origin) and SC-5 (900 meters from origin). At all sampling sites (except for the rocky source SC-1, where no sediment material was observed) it was possible to collect sediment samples: specifically three from the second sampling point (SC-2S1, SC-2S2, SC-2S3), three from the third sampling point (SC-3S1, SC-3S2, SC-3S3), one from the fourth sampling point (SC-4S1) and four from the fifth sampling point (SC-5S1, SC-5S2, SC-5S3, SC-5S4). The temperature, pH and dissolved oxygen (DO) amounts were measured with a dissolved oxygen meter Model 50B (Yellow Springs Instrument Company Inc, Ohio, USA). Water samples for chemical analysis were collected in clean glass bottles, chilled on ice, and stored at 4 °C until analysis. Samples for analysis of microbial communities were collected in clean and sterile glass bottles, and processed within less than 24 h.

**Fig. 2.**
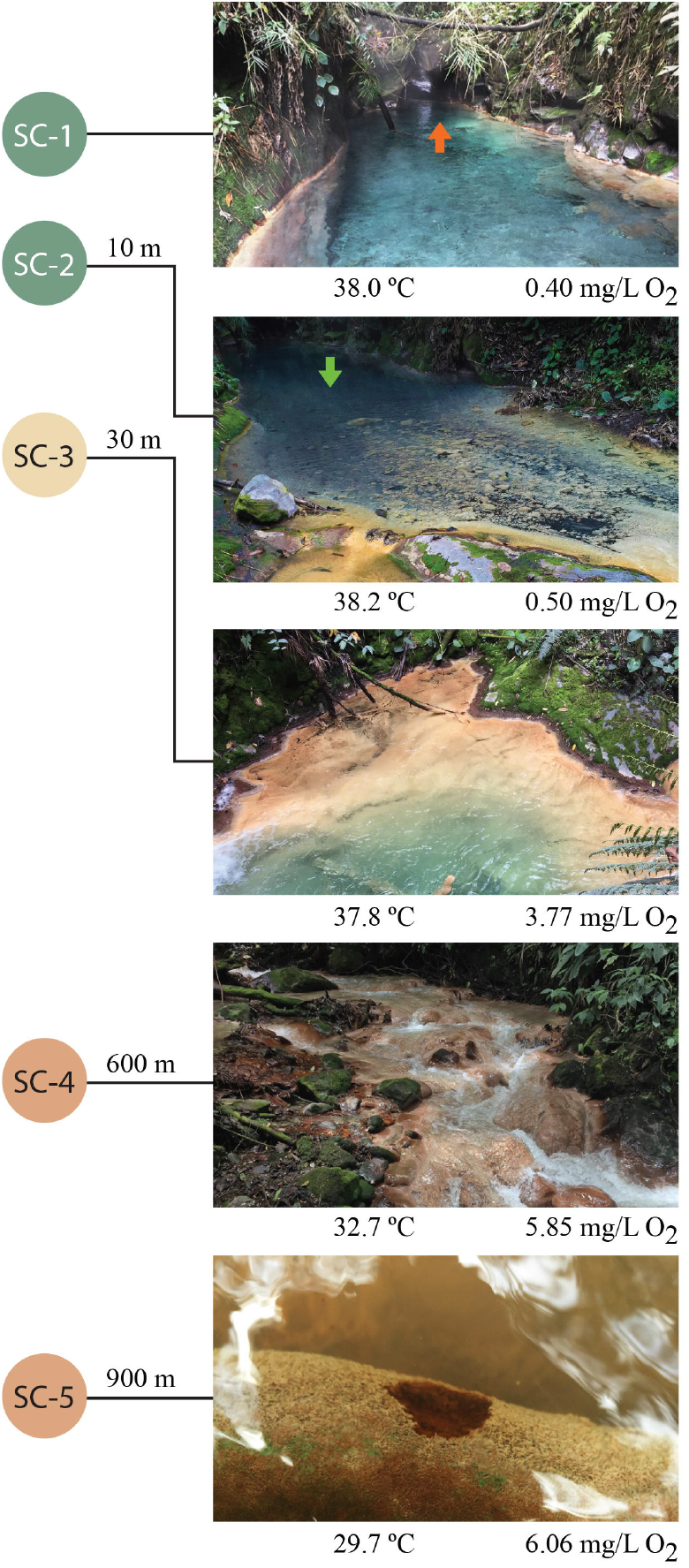
Sampling points from San Cayetano stream. Water and sediment samples were taken in five places along the river: at the origin (SC-1), ten meters from the origin (SC-2), 30 meters from origin (SC-3), 600 meters from origin (SC-4) and 900 meters from origin (SC-5). As shown in the figure, between sampling points a gradient occurs in the oxygen content.

### Chemical Analysis

The water samples were analysed for major anionic and cationic species with an ion-exchange chromatograph (IC, MIC-II, Metrohm Co., Switzerland) and an inductively-coupled-plasma mass spectrometer (ICP-MS, Agilent 7500 instrument, Agilent Technologies, Tokyo, Japan). Samples were filtered with polycarbonate membrane filters (0.45 μm) before analysis. The anions were determined with an IC equipped with an anionic exchange resin (Metrosep A Supp 5 - 100/4.0). Operating conditions were a mobile phase at 33 °C, Na_2_CO_3_ (3.2 mM) / NaHCO_3_ (1.0 mM) and flow rate 0.7 mL/min. The anions were identified and quantified relative to certified commercial standards (Certipur ^®^Anion multi-element standard I, II, Merck, Germany). For ICP-MS analysis a certified multi-element stock solution (Perkin-Elmer Pure Plus standard, product number 9300233) was used. All determinations were performed in triplicates.

### Total DNA isolation, construction of 16S rRNA gene libraries and Illumina sequencing

The three water samples from each sample point (1 L each) were pooled and filtered through a vacuum system under sterile conditions using a membrane filter (pore size 0.22 μm; Millipore, GV CAT No GVWP04700). To prevent rupture, another filter membrane (pore size 0.45 μm; Phenex, Nylon Part No AF0-0504) was placed below. The upper filter was collected and stored at −80 °C until processing. The DNA was extracted from aseptically cut pieces of the filter with a DNA isolation kit (PowerSoil^®^, MoBio, Carlsbad, CA, USA) as described by the manufacturer. Cell lysis was accomplished by two steps of bead beating (FastPrep-24, MP Biomedicals, Santa Ana, CA, USA) for 30 s at 5.5 m s^−1^. To process the sediments, a homogeneous sample of 500 mg was sampled and DNA was extracted using the same protocol. For the construction of microbial 16S rRNA amplicon libraries, the V5-V6 hypervariable regions were PCR-amplified with universal primers 807F and 1050R (Bohorquez *et al*. 2012). The barcoding of the DNA amplicons and the addition of Illumina adaptors were conducted by PCR as described previously (Camarinha-Silva *et al*. 2014). The PCR-generated amplicon libraries were subjected to 250 nt paired-end sequencing on a MiSeq platform (Illumina, San Diego, CA, USA).

### Bioinformatic and phylogenetic analysis of 16S rDNA amplicon data

Bioinformatic processing was performed as previously described (Schulz *et al*. 2018). Raw reads were merged with the Ribosomal Database Project (RDP) assembler (Cole *et al*. 2014), obtaining overall 530,413 paired-end reads. Sequences were aligned within MOTHUR (gotoh algorithm using the SILVA reference database; Schloss *et al*. 2009) and subjected to preclustering (diffs=2) yielding the so-called operational taxonomic units (OTUs) or phylotypes that were filtered for an average abundance of ≥0.001% and a sequence length ≥250 bp before analysis. OTUs were taxonomically classified into the SILVA v132 taxonomy (Yilmaz *et al*. 2014) as reported by the SINA classification tool (Pruesse *et al*. 2012).OTUs were assigned to a taxonomic rank only if their best hit in the SILVA database (Pruesse *et al*. 2007) had an identity higher than the threshold established by Yarza *et al*. (2014) for that rank (94.5% for genus, 86.5% for family, 82.0% for order, 78.5% for class and 75.0% for phylum). Moreover, the sequences of some highly abundant OTUs were also manually examined by means of BLASTN (Altschul *et al*. 1997) against the non-redundant and against the bacterial and archaeal 16S rRNA databases. The statistical analyses and visualizations were performed in R (R Core Team, 2017). We used Vegan (Oksanen *et al*. 2017) to calculate alpha diversity estimators, non-metric multidimensional scaling analyses, (NMDS), and the permutational analysis of variance (Permanova) on normalized tables of OTUs (Supplementary Table S1).

## Results and Discussion

### Physicochemical analysis of San Cayetano stream

The pH along the 0.9 km section of the stream studied in San Cayetano (Fig. 1) was acidic (pH of 3) (Table 1) and the temperature decreased slightly when moving away from the origin (~38° C in SC-1 to ~30° C in SC-5; Fig. 2). Moreover, a prominent gradient in the oxygen concentration was observed throughout the VARD: in SC-1 conditions were almost anoxic (0.40 mg L^−1^ O_2_ corresponding to an oxygen saturation of 6.6% at the temperature and pressure of the sampling site), while at the last sampling point SC-5 water has become oxic (6.06 mg L^−1^ O_2_; 81% O_2_ saturation). As shown in Fig. 2, San Cayetano has its origin in a rocky outcrop within the forest where warm underground water emerges. From there, the water flows for about 3 km to its confluence with the Rio Sucio (Arce-Rodríguez *et al*. 2017). Near the source, very clear water with a depth of about 1.3 m flows smoothly over a clean gravel riverbed, but about 20 m downriver there is a 6 m high waterfall where a light-brown microbial mat is formed. From this point on, water flows faster and it is estimated to discharge at a rate of nearly 100 L/s (Baldoni *et al*. unpublished).

**Table 1.** Physical properties and chemical composition of San Cayetano.

| Property/element/ion | SC-1 | SC-2 | SC-3 | SC-4 | SC-5 |
| --- | --- | --- | --- | --- | --- |
| Temperature (°C) / ± 0.1 | 38.0 | 38.2 | 37.8 | 32.7 | 29.7 |
| pH / ± 0.1 | 3.0 | 2.9 | 3.0 | 3.1 | 3.0 |
| Dissolved oxygen<br>(mg L<sup>-1</sup>) / ± 0.01 | 0.40 | 0.50 | 3.77 | 5.85 | 6.06 |
| Aluminum (mg L<sup>-1</sup>) ± 5 | 41 | 33 | 41 | 34 | 45 |
| Arsenic (µg L<sup>-1</sup>) ± 0.7 | 19.2 | 20.8 | 20.7 | 12.9 | 11.3 |
| Cadmium (µg L<sup>-1</sup>) | < 0.11 | < 0.11 | < 0.11 | < 0.11 | < 0.11 |
| Calcium (mg L<sup>-1</sup>) ± 2 | 4 | 4 | 4 | 4 | 3 |
| Chloride (mg L<sup>-1</sup>) ± 10 | 246 | 203 | 227 | 200 | 222 |
| Copper (mg L<sup>-1</sup>) | < 0.10 | < 0.10 | < 0.10 | < 0.10 | < 0.10 |
| Chromium (µg L<sup>-1</sup>) / ± 2 | 30 | 33 | 33 | 32 | 29 |
| Fluoride (mg L<sup>-1</sup>) | <0.09 | <0.09 | <0.09 | <0.09 | <0.09 |
| Iron (mg L<sup>-1</sup>) / ± 2 | 45 | 51 | 61 | 40 | 35 |
| Magnesium (mg L<sup>-1</sup>) ± 3 | 58 | 64 | 64 | 63 | 59 |
| Manganese (mg L<sup>-1</sup>) ± 0.06 | 1.98 | 2.05 | 1.98 | 2.00 | 2.02 |
| Nickel (µg L<sup>-1</sup>) / ± 2 | 6 | 10 | 8 | 10 | 10 |
| Nitrate (mg L<sup>-1</sup>) | < 0.10 | < 0.10 | < 0.10 | < 0.10 | < 0.10 |
| Lead (µg L<sup>-1</sup>) | < 1.2 | < 1.2 | < 1.2 | < 1.2 | < 1.2 |
| Potassium (mg L<sup>-1</sup>) ± 0.6 | 31.8 | 34.2 | 33.4 | 35.0 | 33.8 |
| Sodium (mg L<sup>-1</sup>) ± 2 | 73 | 79 | 78 | 78 | 75 |
| Sulfate (mg L<sup>-1</sup>) ± 50 | 1190 | 950 | 883 | 870 | 874 |
| Zinc (µg L<sup>-1</sup>) ± 10 | 115 | 109 | 141 | 119 | 121 |

The chemical analysis of filtered samples (Table 1) revealed the presence of sulfate, iron, manganese, aluminium, chromium, zinc, arsenic and magnesium at concentrations much greater than of typical freshwater rivers; the sulfate (~0.87-1.19 g L^−1^) and iron (~35-61 mg L^−1^) levels were particularly high, which is characteristic of volcanic and ARD environments (Arce-Rodríguez *et al*. 2019). The low [iron]/[sulfate] ratio and the absence of precipitated iron minerals (see Figs. 2a,b) as it occurs in Rio Sucio with the iron-oxyhydroxysulfate Schwertmannite (Arce-Rodríguez *et al*. 2017), suggests that much of the sulfate is produced from volcanic sources (*i.e*, from biotic or abiotic H_2_S oxidation and SO_2_ water reaction) rather than the oxidation of pyrites or other metal sulfides. The moderately high temperature (~38°C) that occurs at the origin of San Cayetano also indicates volcanic influence. These observations suggest that the extreme conditions observed in San Cayetano might be produced by discharge products of volcanic or hydrothermal activity into the water bodies of the river, a phenomenon well described in geology ((Vaselli *et al*. 2010). When gases SO_2_ and H_2_S from volcanic origin are intercepted by water bodies on their way to the surface, various chemical reactions might occur: For example, when SO_2_ is discharged into deep underground water bodies this can produce sulfate and acidic conditions (3SO_2_ + 3 H_2_O → 2SO_4_^2-^ + H_2_O + S^0^ + 4H^+^) (Kusakabe *et al*. 2000). On the other hand, if present, H_2_S may be subjected to biotic or abiotic oxidation to also generate sulfate (e.g. H_2_S + 2 O_2_ → H_2_SO_4_). In any case, it is clear that the low [iron]/[sulfate] ratio, reflects that there is a sulfate source other than the dissolution and oxidation of minerals such as pyrite, i.e. from volcanic origin. For example, in Río Tinto ARD system the [Fe]/[SO_4_^2-^] molar ratio is 0.58 at the origin and only decreases to 0.29 downstream as expected from the 0.5 ratio from oxidation of pyrite FeS_2_; however, in San Cayetano stream, we measure far lower ratios between 0.07 at the origin and 0.07-0.12 downstream. Our ratios compare well to the 0.15 and 0.06 ratios at Copahue Volcano’s (Urbieta *et al*. 2015).

Both processes (ARD or volcanic activity) produce acid that favors the dissolution of minerals (e.g. iron oxides, sulfides and aluminosilicate rocks). This correlates well with the acidic pH of San Cayetano (pH 2.94-3.06) and the presence of dissolved metals such as aluminum, zinc, arsenic, manganese or chromium. In summary, the physicochemical parameters suggest that: (i) the selected sampling points describe a gradient of increasing oxygen concentration starting at the origin with almost anoxic conditions and (ii) San Cayetano has the typical chemical composition of an ARD environment but with an important contribution of sulfuric acid due to volcanic processes. For that reason, we have proposed the acronym VARD (volcanic influenced acid rock drainage) to describe this specific type of ARD ecosystem.

### The concentrations of dissolved oxygen along the river shapes the microbial community of San Cayetano

In spite of the acidic conditions and high concentrations of sulfate, iron and toxic metals, a relatively high diversity of microorganisms was observed in the San Cayetano VARD. We identified in total 1904 OTUs (excluding singletons) from 530,310 16S rRNA gene sequences originating from 36 phyla and 177 families from both bacteria and archaea (Fig. 3 and Supplementary Table 1). According to the relative percentage of sequences, the most abundant phylogenetic groups were Gammaproteobacteria (40.2%), Euryarchaeota (10.0%), Actinobacteria (9.1%), Nitrospirae (6.4%) and Firmicutes (6.2%). However, we detected important variations between the different sampling sites (Fig. 3). These results are consistent with the alpha-diversity estimations showing, for example, that 63% of the samples in San Cayetano presented richness values higher than 500 OTUs, and that 88% of the samples presented values of the Shannon index greater than 3 (Supplementary Figure S1).

Despite the high abundance of microbes found in San Cayetano, the taxonomic composition of this VARD showed a remarkable abundance of unclassified microorganisms amongst the most represented OTUs, both in water and sediment samples. This was specially the case for the water samples taken at the first two sampling points, where the dissolved oxygen concentrations do not exceed 0.50 mg L^−1^. Nevertheless, we could detect the presence of microorganisms commonly found in ARD environments like iron-, sulfur-, sulfide- and thiosulfate-oxidizing bacteria. Furthermore, we determined significant differences in the community structure at the OTU level (Permanova, P=0.001) along the oxygen gradient.

NMDS and Permanova analyses were consistent at showing the role of oxygen availability in shaping the structure of the microbial communities in the VARD ecosystem (Fig. 4). We observed a clear separation of the samples according to the habitat within each oxygen condition. This difference between microbial communities in the water column respect to those in sediments was significant (Permanova, P=0.018). The water samples near the origin (SC-1 and SC-2) that contain very little concentrations of oxygen reflect a very similar microbial community amongst each other. The same situation was observed among the sediments obtained in the second sampling point (SC-2S1, SC-2S2 and SC-2S3). As expected, samples from the third sampling point, containing intermediate levels of oxygen, are not grouped with SC-1 and SC-2 (low oxygen) nor with SC-4 and SC-5 (high oxygen). Again the sediments at this point (SC-3S1, SC-3S2 and SC-3S3) are grouped together. Finally, the water samples of points four and five, as well as their respective sediments are clearly grouped. As already mentioned, the last two sampling points present very similar aerobic conditions, so it is expected that the microbial community in these sites will be grouped as shown by the NMDS analysis.

**Fig. 3.**
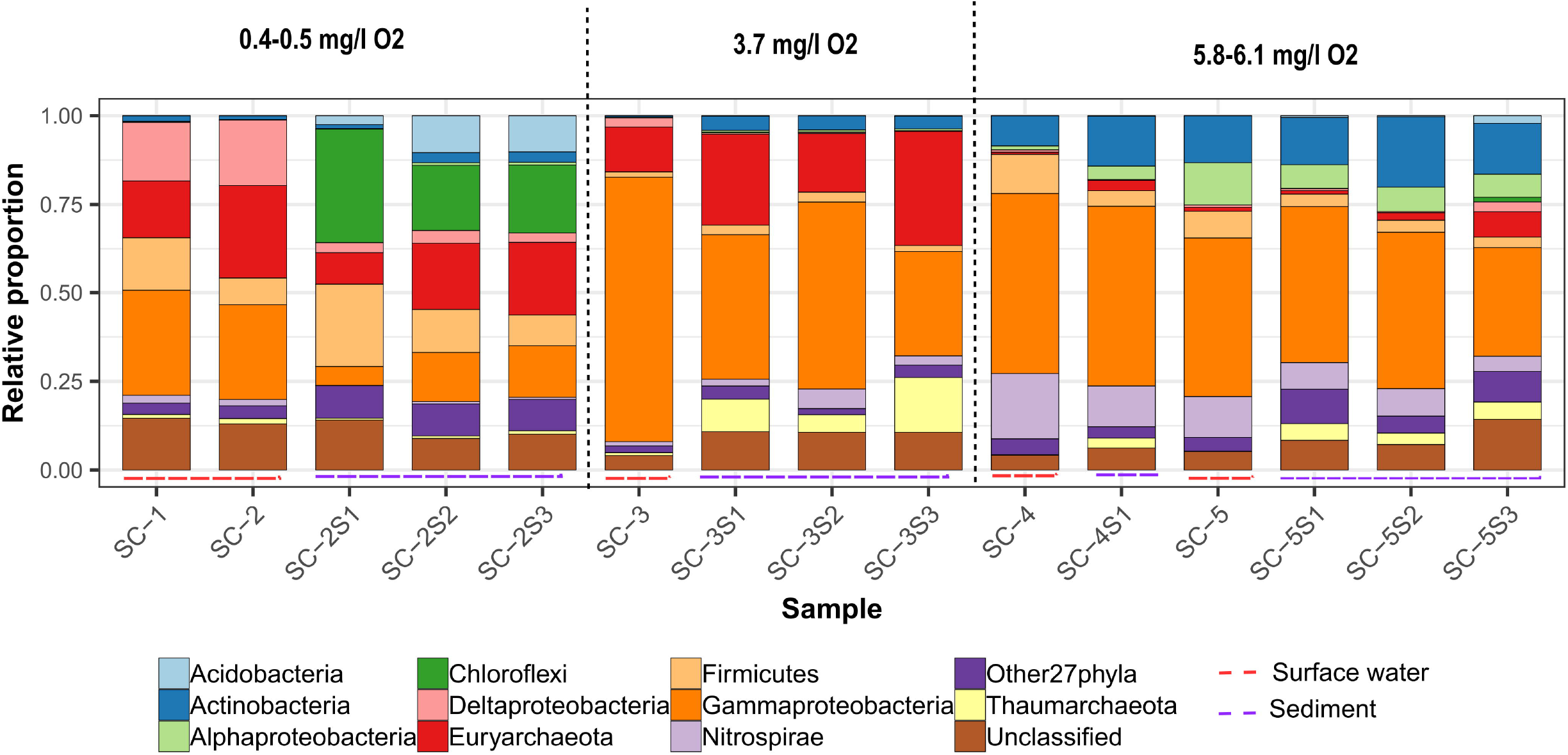
Taxonomic composition of San Cayetano. Relative abundance of bacterial and archaeal organisms to the phylum level. The OTUs were taxonomically classified into the RDP taxonomy as reported by the SINA classification tool, as described in Materials and Methods. The water samples at each of the sampling points are identified as SC-1 to SC-5. Sediment samples are identified with legends SC-2S1 to SC-5S4.

As mentioned above, in the first sampling point (SC-1), which corresponds to the spring, the waters contain only 0.40 mg L^−1^ O_2_. At this site, the water springs from the subsurface between the rocks. The microbial community of the water sample (SC-1 in Fig. 5 and Supplementary Table 1) is dominated by an unclassified Deltaproteobacterium (OTU_SC0012; 14.8% of the total reads in SC-1), with high (97.2%) similarity to environmental OTUs obtained from other AMD/ARD environments such as Río Tinto, Spain (NCBI accession FN862110) or the TongLing acid mine drainage and the Dexing Copper Mine in China (NCBI accessions KC749174 and EF409870, respectively. Interestingly, this Deltaproteobacterium was also similar (96.8%) to the uncultured phylotype K5_62 (NCBI accession EF464599), which has been proposed to represent a novel type of acidophilic sulfate-reducing bacteria (Winch *et al*. 2009). Given the high concentrations of sulfate that exist in the origin of San Cayetano (~1.19 g L^−1^), the presence of microorganisms capable of using this oxyanion as the terminal electron acceptor is expected. The second most abundant microorganism in the origin of San Cayetano (OTU_SC0019; 14.5%) corresponds to an unclassified bacterium according to our classification methodology. A closer look into its sequence by means of BLAST revealed that the first 183 nucleotides of the DNA fragment share 96.7% of similarity with *Sulfobacillus thermosulfidooxidans*, while the rest of the sequence corresponds to a low complexity, GC-rich DNA region that does not match with any other sequence within the non-redundant nucleotide collection (Supplementary Table S1). We, therefore, resolved to manually curate the sequence of this this OTU and we reassign it within genus *Sulfobacillus* (Fig. 5). These bacteria are Gram-positive, moderately thermophilic, facultative anaerobic (Johnson *et al*. 2008) acidophiles that grow in sulfurous and sulfidic environments. It has been previously reported that they attach to sulfide mineral surfaces, which may promote faster sulfide oxidation (Watling, Perrot and Shiers 2008). *Sulfobacillus* species are also involved in ferrous-iron oxidation (Norris *et al*. 1996; Pina *et al*. 2010). In addition to these two highly represented genera at the origin of San Cayetano, we identified other microorganisms related to acidophilic, sulfur- and iron-oxidizing bacteria, such as *Acidithiobacillus* (OTU_SC0016; 12.5%). This genus includes chemolithotrophic bacteria capable of oxidizing sulfide to sulfate, coupling this reaction to ferric iron (under anoxic conditions) or oxygen (under oxic conditions) reduction (Suzuki *et al*. 1990; Pronk and Johnson 1992). Together with *Acidithiobacillus*, another putative chemolithotrophic sulfur oxidizer from family Hydrogenophilaceae was found in sample SC-1 (OTU_SC0015; 8.9%). This organism is closely related to a group of bacteria capable to obtain energy by the oxidation of sulfur compounds with oxygen or nitrate as the terminal electron acceptor, including *Sulfuritortus calidifontis* (93.2% of similarity according to BLAST), *Annwoodia aquaesulis* (92.8% similarity) and *Thiobacillus thioparus* (92.4% similarity)(Kojima, Watanabe and Fukui 2017; Boden, Hutt and Rae 2017).

**Fig. 4.**
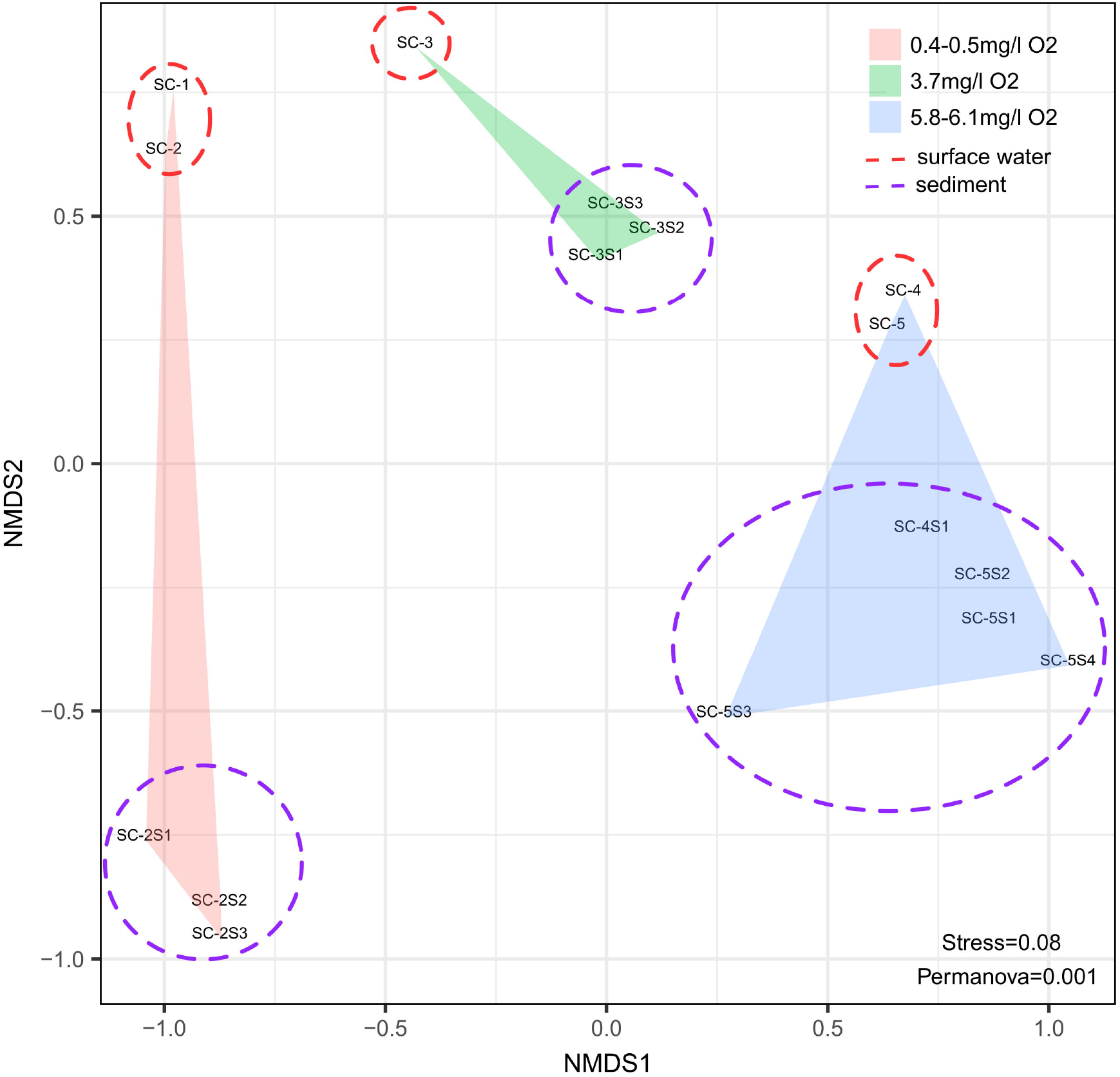
Non-metric multidimensional scaling analysis of the prokaryotic communities in San Cayetano volcanic influenced acid rock drainage. A clustering of communities according to the oxygen content of the samples and also by the habitat (water column versus sediment) is shown. The NMDS and the Permanova analyses were performed with the package Vegan.

**Fig. 5.**
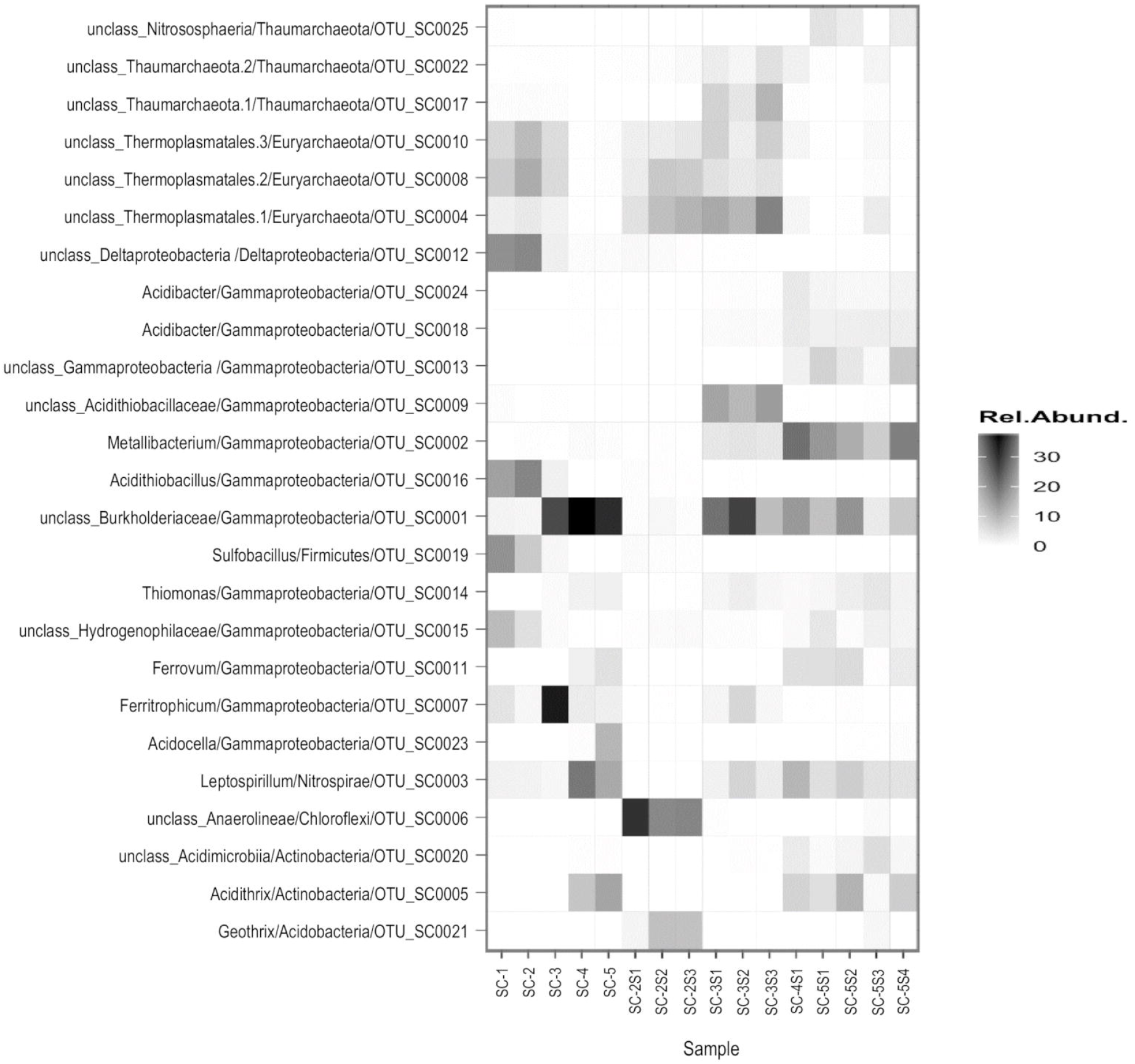
Heat map representing the most abundant genera in each sample. The heat map depicts the relative percentage of 16S rRNA gene sequences assigned to each genus (y axis) across the 16 samples analysed (x axis). Square colors shifted towards black indicate higher abundance.

Two archaea from the order Thermoplasmatales were also detected amongst the most abundant microorganisms in the water samples from SC-1 (OTU_SC0008; 6.6% and OTU_SC0010; 5.0% of total SC-1 reads). Both phylotypes share high similarity to environmental OTUs detected in other AMD/ARD niches such as the Rio Tinto (NCBI accession FN862291), the hydrothermal ponds in the Copahue region in Argentina (NCBI accession JX989254) or the Kamchatkan hot springs in Russia (NCBI accession JF317816). Moreover, the closest isolates in both cases belong to genus *Cuniculiplasma* (Fig. 5 and Supplementary Table 1). Archaea from the order Thermoplasmatales are considered as extreme thermoacidophiles that generally grow at pH less than 4 (Auernik, Cooper and Kelly 2008). Specifically, *Cuniculiplasma* is classified as a facultative anaerobic, organotrophic and mesophilic whose only isolate (*C. divulgatum*) was obtained from acidic streamers formed on the surfaces of copper-ore-containing sulfidic deposits in south-west Spain and North Wales, UK (Golyshina *et al*. 2016). Other species of order Thermoplasmatales such as *Thermoplasma acidophilum* (Yasuda *et al*. 1995), *Thermoplasma volcanium* (Segerer *et al*. 1988), *Thermogymnomonas acidicola* (Itoh, Yoshikawa and Takashina 2007), and *Picrophilus torridus* (Schleper *et al*. 1995; Serour and Antranikian 2002) have been obtained from solfataric hydrothermal areas or acidic streamers containing sulfidic deposits, i.e. sulfur-rich environments. Thus, it is likely that the two Thermoplasmatales OTUs found at the spring waters of San Cayetano are also involved in sulfur metabolism.

In the second sampling point (SC-2), just 10 meters from the origin, the content of dissolved oxygen only undergoes a slight increase (0.50 mg L^−1^ O_2_, 7.7% O_2_) so the microbial community does not change drastically. In this site, the microorganisms found in SC-1, i.e., *Acidithiobacillus* (16.2%), the putative sulfate reducing Deltaproteobacterium (15.9%), both Thermoplasmatales archaea (OTU_SC0008; 10.9% and OTU_SC0010; 8.9%), and the *Sulfobacillus* (6.9%), are maintained in a major proportion. At this sampling point, it was possible to assess also the microbiota in the sediments. The most abundant microorganism in all the sediment samples at SC-2 is an unclassified Anaerolineae chloroflexi bacterium (OTU_SC0006) distantly related to genera *Anaerolinea* (89.8% similar to *Anaerolinea thermolimosa), Ornatilinea* (88.0% similar to *Ornatilinea apprima*) and *Caldilinea* (85.5% similar to *Caldilinea aerophila*). These genera are reported as chemo-organoheterotrophic chloroflexi, which thrive in very diverse habitats such as hot spring aquifers (Grégoire *et al*. 2011) or anaerobic wastewater sludge (Sekiguchi *et al*. 2003; Yamada *et al*. 2005). Interestingly, we could also identify the presence of *Geothrix* in the sediments of sample SC-2 (OTU_ SC0021). Only one species from this genus, *Geothrix fermentans*, has been isolated to date. It is a strict anaerobe capable to oxidize a wide variety of organic acids with ferric iron serving as the sole electron acceptor (Coates *et al*. 1999). This organism could play a very important role in the sediments of the San Cayetano ecosystem, not only by the oxidation of organic matter but also in the dissolution of iron(III). By doing so, the iron(II) produced by its respiration becomes available for other iron-oxidizing bacteria along the river course (i.e. *Ferritrophicum* spp. *Leptospirillum* spp., *Acidithrix* spp., *Ferrovum* spp., see below).

Besides one of the Thermoplasmatales microorganism found also in the water sample (OTU_SC0008), another archaeon of the same order appeared as highly abundant in the sediments of SC-2 (OTU_SC0004; Fig. 5). Albeit it also belongs to an unclassified species, it shows high (99.2%) similarity to environmental OTUs obtained from the Kamchatkan hot springs (NCBI accession JF317816) or the sulfuric ponds of the Tatung Volcano area in Taiwan (NCBI accession FJ797335). The closest isolates from this OTU lie within the *Picrophilus* genus (90.7% similarity to both *P. torridus* and *P. oshimae* (Schleper *et al*. 1995). These microorganisms represent the most thermoacidophilic organisms known, with the ability to survive at a pH around 0 (Schleper *et al*. 1995; Fütterer *et al*. 2004). Strains of this species were first isolated from a dry solfataric field in Japan (Fütterer *et al*. 2004). Thus, it is possible that both Thermoplasmatales archaea take some part in the metabolism of sulfur in the sediments at this sampling point.

At the third sampling point (30 meters from the source, after a small waterfall) there is a significant change in the oxygen content (3.77 mg L^−1^ O_2_; 58% O_2_) as well as in the microbial community. Interestingly, more than 55% of the microbial community in the water sample (SC-3) is dominated by two Gammaproteobacteria: a bacterium of genus *Ferritrophicum* (OTU_SC0007; 33.2%) characterized for metabolizing iron compounds, and an unclassified Burkholderiaceae (OTU_SC0001; 25.1%)(Fig. 5). Of the genus *Ferritrophicum* only one species has been reported, *F. radicicola*, which uses ferrous iron as an energy source for lithotrophic growth. It is a microaerophilic bacterium isolated from a plant growing in an acid mine drainage system (Weiss *et al*. 2007). Thus, it is highly possible that the *Ferritrophicum* from San Cayetano participates in the oxidation of dissolved ferrous iron. On the other hand, representative species of Burkholderiaceae family are very abundant, occupying diverse ecological niches and performing very diverse metabolic functions (e.g. nitrogen fixation, aromatic compounds catabolism, etc) (Compant *et al*. 2008). Therefore, it is difficult to assign a specific role to this generalist Betaproteobacterium in San Cayetano VARD. The *Acidithiobacillus* and the two Thermoplasmatales found in the waters of SC-2 were also present in this sample in lower abundance.

The sediments of SC-3 also show the presence of the same Burkholderiaceae bacterium (OTU_SC0001) found in the water sample, as well as the Thermoplasmatales archaea (OTU_SC0004 and OTU_SC0008, respectively) found in the sediments of SC-2 (Fig. 5). The presence of the two archaea suggests their capacity to adapt to a wide range of oxygen conditions. In addition, we found a highly abundant Gammaproteobacterium that clusters within the Acidithiobacillaceae group RCP1-48 (OTU_SC0009). Members from this group have been previously detected in the sediments of other AMD habitats like Rio Tinto (Sánchez-Andrea *et al*. 2011) or Los Rueldos mercury underground mine in Spain (Mesa *et al*. 2017), and it has been suggested that they could have an important role in the oxidation of iron and sulfur compounds (Mesa *et al*. 2017). Besides those microorganisms, these sediments host two other highly abundant iron oxidizing-bacteria: A representative species of genus *Leptospirillum* (OTU_SC0003), and the same *Ferritrophicum* sp. found in the water sample. Particularly, the presence of *Leptospirillum* has been widely reported in acid mine drainage (AMD) environments. They catalyze the ferrous iron oxidation, accelerating iron(III)-mediated oxidative dissolution of sulfide minerals and thus the formation of AMD (Goltsman *et al*. 2013). Due to its extraordinary characteristics, *Leptospirillum* species have been used throughout the world in industrial bioleaching operations (Sand *et al*. 1992; Issotta *et al*. 2016). In summary, at sampling point SC-3 we observe an important increase of heterotrophic and iron-oxidizing bacteria, while anaerobic sulfur oxidizers drastically decrease (compared to samples SC-1 and SC-2). This observation also correlates with the increase of oxygen concentration at SC-3 (see table 1 and Fig. 4).

Finally, the sampling points four and five (600 m and 900 m from the origin, respectively) have similar high oxygen contents and very similar microbial composition (Figs 4 and 5). The oxygen content was 5.85 mg L^−1^ O_2_ (82% O_2_) and 6.06 mg L^−1^ O_2_ (81% O_2_) respectively. The microbial community in those waters is also dominated by the same Burkholderiaceae bacterium found in SC-3 (36.9% in SC-4 and 30.0% in SC-5), followed by the *Leptospirillum* sp. present in the sediments of SC-3 (18.3% in SC-4 and 11.4% in SC-5). In the third place, we detected large quantities of the heterotrophic acidophile *Acidithrix* (OTU_SC0005; 7.58% in SC-4 and 11.88% in SC-5). To the best of our knowledge, *Acidithrix ferrooxidans* is the unique species from this genus that has been described to date. This bacterium is capable to catalyze the dissimilatory oxidation of ferrous iron under aerobic conditions, and the reduction of ferric iron under micro-aerobic and anaerobic conditions (Jones and Johnson 2015). Given the oxic nature of SC-4 and SC-5, it is presumable that the *Acidithrix* detected in both water samples contributes to the oxidation of ferrous iron. These conditions are very appropriate also for the obligate aerobe *Leptospirillum* sp. (see above) and for other iron-oxidizing bacteria present in these samples in minor abundance (i.e *Ferritrophicum* spp. and *Ferrovum* spp.). In addition, other heterotrophic bacteria from the genera *Trichococcus* and *Acidocella* were found in the water from sampling points SC-4 and SC-5, respectively.

Finally, the sediments obtained from SC-4 and SC-5 showed a similar microbial composition (Fig. 5). The most prominent organism in both samples is a Gammaproteobacterium of the genus *Metallibacterium* (OTU_SC0002). Once again, this genus has only one reported species (*M. scheffleri*) and has been very poorly studied. Among the few known metabolic functions of this species, it is known that it is capable of reducing ferric iron, but does not oxidize ferrous iron (Ziegler *et al*. 2013). We found also a large abundance of some of the microorganisms identified in the water samples from sampling points SC-4 and SC-5, such as the Burkholderiaceae Betaproteobacterium, *Acidithrix sp., Leptospirillum* sp. and *Ferrovum* sp. Other microorganisms found in the sediments of SC-5 were a Gammaproteobacterium (OTU_SC0013) from the uncultured group KF-JG30-C25 and another Actinobacterium (OTU_SC0020) whose closest isolate (91.4% similarity according to BLAST) is the heterotrophic iron(III)-reducer *Aciditerrimonas ferrireducens* (Itoh *et al*. 2011). Specifically, members of KF-JG30-C25 group have been detected in AMD habitats such as the uranium mining waste pile at Johanngeorgenstadt in Germany (Selenska-Pobell 2002).

## Conclusions

This work reports the microbiological composition of the volcanic influenced acid rock drainage known as San Cayetano (northeast of Irazu volcano, Costa Rica). The low [iron]/[sulfate] ratio suggest that most of the sulfuric acid in this VARD is of volcanic origin, possibly due to the disproportionation reaction of SO_2_ and biotic or abiotic H_2_S oxidation in the groundwater. The moderately high temperature of the water at the spring of San Cayetano is also consistent with underground mixing with volcanic water. We analyzed the microbial composition of water and sediment samples along 900 meters from the origin of the river. Interestingly, we found a remarkable abundance of a yet not-described diversity of microorganisms amongst the most represented taxa thriving in San Cayetano VARD. Nevertheless, we could also identify the presence of bacteria and archaea that, along with the chemistry of the site, resembles the composition of typical ARD environments. The structure of the microbial community in San Cayetano undergoes important changes as a function of the oxygen gradient observed in the distinct water samples. At the origin of the river (SC-1 and SC-2), the microbial profile is dominated mainly by putative anaerobic, sulfur-oxidizing microorganisms such as *Sulfobacillus* sp., *Acidithiobacillus* sp., the putative sulfate reducing Deltaproteobacterium and the *Hydrogenophilaceae* Gammaproteobacterium. Their presence at the first two sampling points, suggest that these microorganisms are likely involved in both sulfate reduction as well sulfide oxidation, completing the sulfur cycle. At the third sampling point (SC-3), the oxygen content increases eight-fold (from 0.40 mg L^−1^ to 3.77 mg L^−1^), also generating a drastic change in the microbial community. Most of the sulfur-oxidizing taxa found at the origin of the river are substituted by iron-oxidizers (*Ferritrophicum spp., Leptospirillum sp.*) and heterotrophic microorganisms (Burkholderiaceae Betaproteobacterium). After 600 meters downstream from the origin of San Cayetano, the river is completely oxygenated. Consequently, the last two samples (SC-4 and SC-5) are totally dominated by aerobic microorganisms, mostly related to heterotrophic metabolism (Burkholderiaceae Betaproteobacterium, *Trichococcus* sp. *Acidocella* sp.) or iron-oxidation (*Leptospirillum* sp., *Acidithrix* sp., *Ferritrophicum* spp., *Ferrovum* spp.). Our data are consistent with the notion that the initial volcanic chemistry of the aquifer is fundamentally modified by the microbial metabolism and by the increase of oxygen along the river course until its confluence with the previously characterized Rio Sucio (Arce-Rodriguez *et al*. 2017).

Over the 900 meters course of the stream analyzed, we were able to find both aerobic and anaerobic organisms carrying out many different metabolic activities (*i.e*. sulfur- and iron- oxidation/reduction, nitrogen metabolism, carbon oxidation, etc.). Furthermore, these organisms have to cope with the high concentration of heavy metals such as arsenic, chromium or zinc. Taking also into account the large diversity of unclassified microorganisms found along San Cayetano, it is reasonable to presume that this untapped ecosystem is endowed with metabolic functions and enzymatic activities with a biotechnological interest that could be further exploited. This work also contributes to the knowledge of volcanic and ARD environments in which the conditions are created only by natural factors which contrasts with the majority of published works on ARD environments in the world which have been influenced by human activity.

## Supporting information

Table S1. DNA sequence and phylogenetic assignment of the most abundant phylotypes detected in San Cayetano.

Figure S1. Alpha-diversity estimations of San Cayetano samples.

## Funding

This work was supported by The Vice-rectory of Research of Universidad de Costa Rica (project number VI 809-B6-524), the Costa Rican Ministry of Science, Technology and Telecommunication (MICITT) and Federal Ministry of Education and Research (BMBF) (project VolcanZyme contract No FI-255B-17) and by the ERC grant IPBSL (ERC250350-IPBSL). F.P-S. is supported by the Spanish Economy and Competitiveness Ministry (MINECO) grant CTM2016-80095-C2-1-R.

## Acknowledgements

We thank Carlos Rodriguez of Centro de Investigación en Contaminación Ambiental (CICA) for the help in the chemical analysis. We also are grateful to Solange Voysest for help in the design of some figures.

