## Supplementary information for "Untapped microbial composition along a horizontal oxygen gradient in a Costa Rican volcanic influenced acid rock drainage system"

^1^Microbial Interactions and Processes Research Group, Helmholtz Centre for Infection Research, 38124, Braunschweig, Germany ^2^Systems Biology Program, Centro Nacional de Biotecnología (CNB-CSIC), C/Darwin 3, 28049 Madrid, Spain ^3^Centro Nacional de Innovaciones Biotecnológicas (CENIBiot), CeNAT-CONARE, 1174-1200, San José, Costa Rica ^4^Escuela de Química, Universidad de Costa Rica, 11501-2060, San José, Costa Rica ^5^Escuela Centroamericana de Geología, Universidad de Costa Rica, 11501-2060, San José, Costa Rica ^6^Laboratorio de Ecología Urbana, Universidad Estatal a Distancia, 11501-2060, San José, Costa Rica ^7^Escuela de Biología, Universidad de Costa Rica, 11501-2060, San José, Costa Rica ^8^Centro de Investigaciones en Productos Naturales (CIPRONA), Universidad de Costa Rica, 11501-2060, San José, Costa Rica.

***Keywords:*** Costa Rica, San Cayetano, Acid Rock Drainage, Microbial Communities, Oxygen gradient*.*

**Table S1.** DNA sequence and phylogenetic assignment of the most abundant phylotypes detected in San Cayetano using Illumina-based amplicon deep-sequencing.

**See Excel file.**

**LEGENDS OF FIGURES**

**Figure S1. Alpha-diversity estimations of San Cayetano samples**. Richness indicate that 63% of the samples presented richness values higher than 500 OTUs, and that 88% of the samples presented values of the Shannon index greater than 3.
