## Supplementary figures and images for "Untapped microbial composition along a horizontal oxygen gradient in a Costa Rican volcanic influenced acid rock drainage system"

### Figure S1. Alpha-diversity estimations of San Cayetano samples.

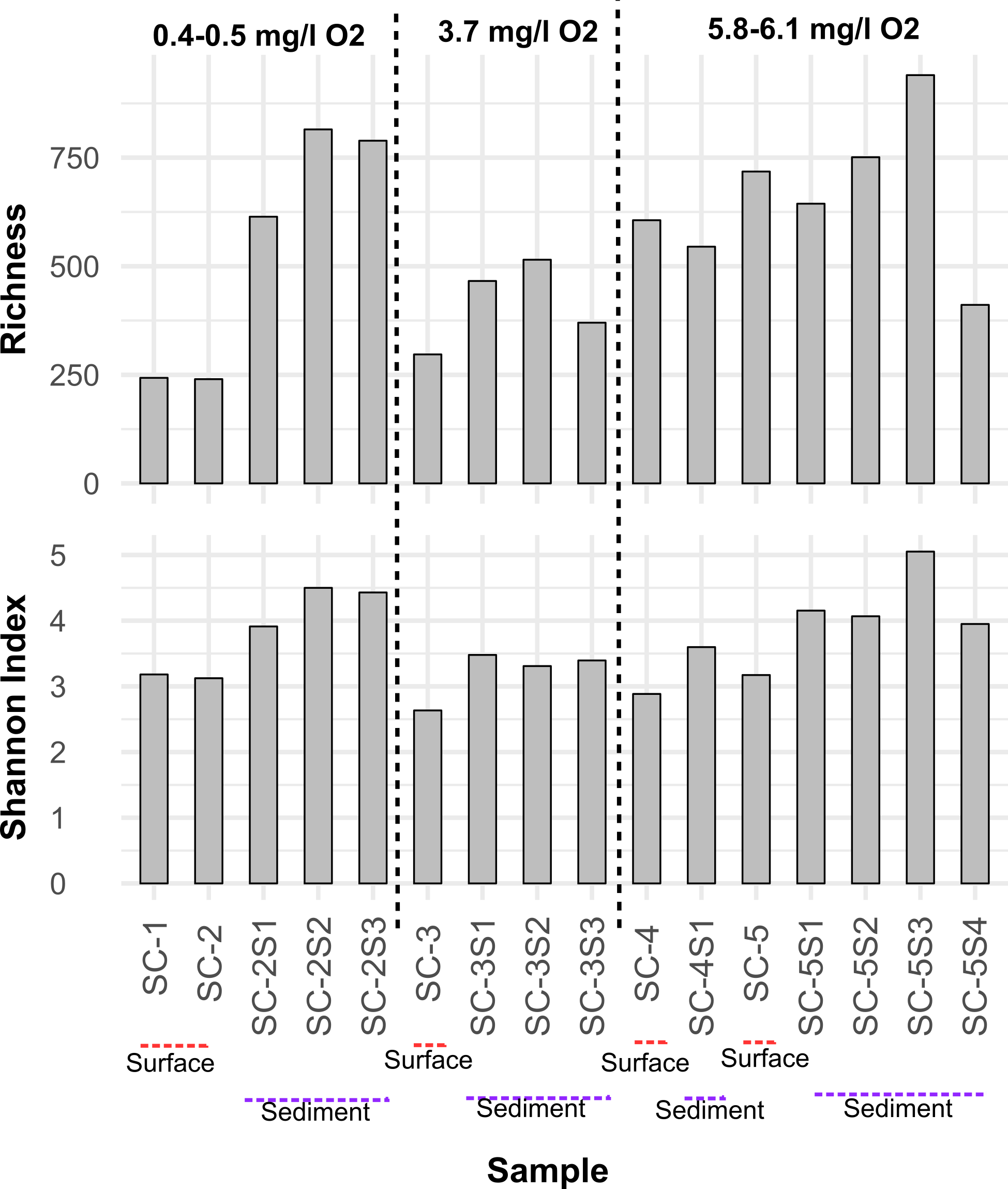
